# EnsembleFlex: Protein Structure Ensemble Analysis Made Easy

**DOI:** 10.1101/2024.12.21.629432

**Authors:** Melanie Schneider, José Antonio Marquez, Andrew R Leach

## Abstract

EnsembleFlex is a novel tool designed to perform dual-scale flexibility analysis of protein structure ensembles, encompassing both backbone and side-chain dynamics. It integrates dedicated superposition methodologies, two dimension reduction techniques, customizable clustering, an automated ligand binding site analysis and analysis of conserved water molecules, enhancing the accuracy of ensemble flexibility analysis and visualisation. With a user-friendly, no-coding graphical interface as well as a code-based option, EnsembleFlex is accessible to a broad range of researchers. The tool complements experimental ensemble data analysis with predictive methods based on elastic network models, bridging the gap between experimental and computational approaches in protein flexibility analysis, and does so from a global backbone-based perspective, as well as from a focused side-chain and binding site-centred point of view. The need for reliable, streamlined, fast, and user-friendly tools for structural ensemble analysis is critical in today’s research environment, and EnsembleFlex addresses these needs efficiently, offering an accessible solution that integrates powerful analysis methods in a single package. Its wide applicability across various domains of molecular design underscores the importance of streamlined flexibility analysis in scientific research.

## Introduction

### Background

Protein flexibility is central to understanding protein function, stability, and interactions. While traditional structural biology has provided critical insights into static protein structures, the dynamic behaviour of proteins is increasingly recognized as essential for their biological activity. This dynamic nature encompasses various scales, from local fluctuations of side chains to large-scale domain movements, crucial in processes such as enzymatic catalysis, molecular recognition, and signal transduction.^1–3^ Understanding protein structural flexibility is not merely a molecular curiosity; it is key to deciphering diseases, designing targeted therapies, and engineering proteins with novel functionalities. ^4–6^

### Current Tools and Gaps: An overview of protein flexibility assessment methods

#### From using uniform computationally generated ensembles to heterogeneous experimentally generated ensembles in the age of big data in structural biology

Several tools exist for computationally “predicting” protein flexibility starting from a single structure by first generating an ensemble and subsequently analysing it. The standard method for generating an ensemble, which is the most widely used and most physics based, is molecular dynamics (MD), including enhanced sampling methods.^7,8^ The main limitation here is that those MD based methods are computationally expensive and not always straightforward to set up. Other methods, thus, take a more coarse-grained and faster approach, particularly through normal mode analysis (NMA) based on elastic network models that primarily focus on backbone flexibility, often neglecting the significant contributions of side-chain dynamics due to higher computational cost.^9–11^ On the other hand the increased level of automation in protein structure determination ^12,13^ has lead to an increase in the availability of experimental structures. Currently, the wwPDB contains multiple examples of highly redundant datasets where the same molecular structure has been independently determined multiple times and where conformational differences, potentially relevant for protein dynamics, can be observed. However, there is a critical gap in tools that can reliably analyse both backbone and all-atom dynamics in a fast, streamlined, and user-friendly manner, especially when dealing with experimentally generated heterogeneous structural ensembles (where atoms and/or residues differ across the ensemble). EnsembleFlex is designed to address this gap by offering an integrated solution that allows researchers to perform complex flexibility analyses without needing extensive coding knowledge. This is becoming increasingly important with the growing data in structural biology.

#### Further current limitations

Additionally, most existing tools require coding expertise, limiting accessibility. Moreover, standard superposition methods used in these tools might not be optimal for flexibility analysis, potentially leading to inaccurate results.^4,14^ There is also a lack of integration between data-driven analyses and predictive models, which can provide complementary insights.^15–17^

### Objective

EnsembleFlex addresses these gaps by providing a comprehensive tool that analyses both backbone and all-atom flexibility of structure ensembles in a user-friendly environment. The dual-scale flexibility analysis is one of the key innovations of EnsembleFlex, as it allows researchers to capture both the large-scale backbone movements and the finer details of side-chain flexibility. These two layers of information combine to give a complete picture of protein dynamics. It offers a robust superposition method tailored for flexibility analysis, two different dimension reduction techniques, customisable clustering, an automated ligand binding site analysis and a conserved water analysis. This holistic approach is particularly important in applications such as drug design, where both backbone movements and side-chain dynamics at the binding site can influence ligand binding and protein functionality.

EnsembleFlex integrates both data-driven and predictive approaches to deliver a more accurate and holistic understanding of protein flexibility.

EnsembleFlex can also be seen as a service extension of the High Throughput Crystallization and Fragment Screening Facility (HTX lab) at EMBL Grenoble, supported by EMBL-EBI. It aims to provide users and the wider scientific community with streamlined tools for the analysis of structural ensembles generated during high-throughput experiments, ensuring data accessibility and usability in downstream applications.

## Methods

### Tool Architecture

EnsembleFlex is built on a modular architecture that integrates computational methods for superpositioning, flexibility analysis (including dimension reduction and clustering), automated ligand binding site analysis, conserved water analysis, and predictive methods within a consistent and user-friendly Python and R-based framework. Implemented functions come from open source R libraries and Python packages with permissive licences. ^4,18–24^ Users can access the tool through a browser-based graphical user interface (GUI), powered by Streamlit ^25^, or via command line scripts, allowing for flexibility in usage. The GUI offers a no-coding option, while the command line provides a fully customizable environment. The tool can be installed via Docker, ensuring reproducibility across systems, or through Conda for system-specific setups. Since EnsembleFlex operates locally without requiring a server connection, all data remains on the user’s system, ensuring complete privacy and eliminating any data-sharing concerns. Default settings for the standard workflow allow beginners to rapidly execute analyses and retrieve results, without extensive parameter customization or in-depth knowledge about underlying functions. An overview of the provided analysis workflow is sketched in Figure 1 and a GUI screenshot is available in supplementary Figure S1.

**Fig. 1:**
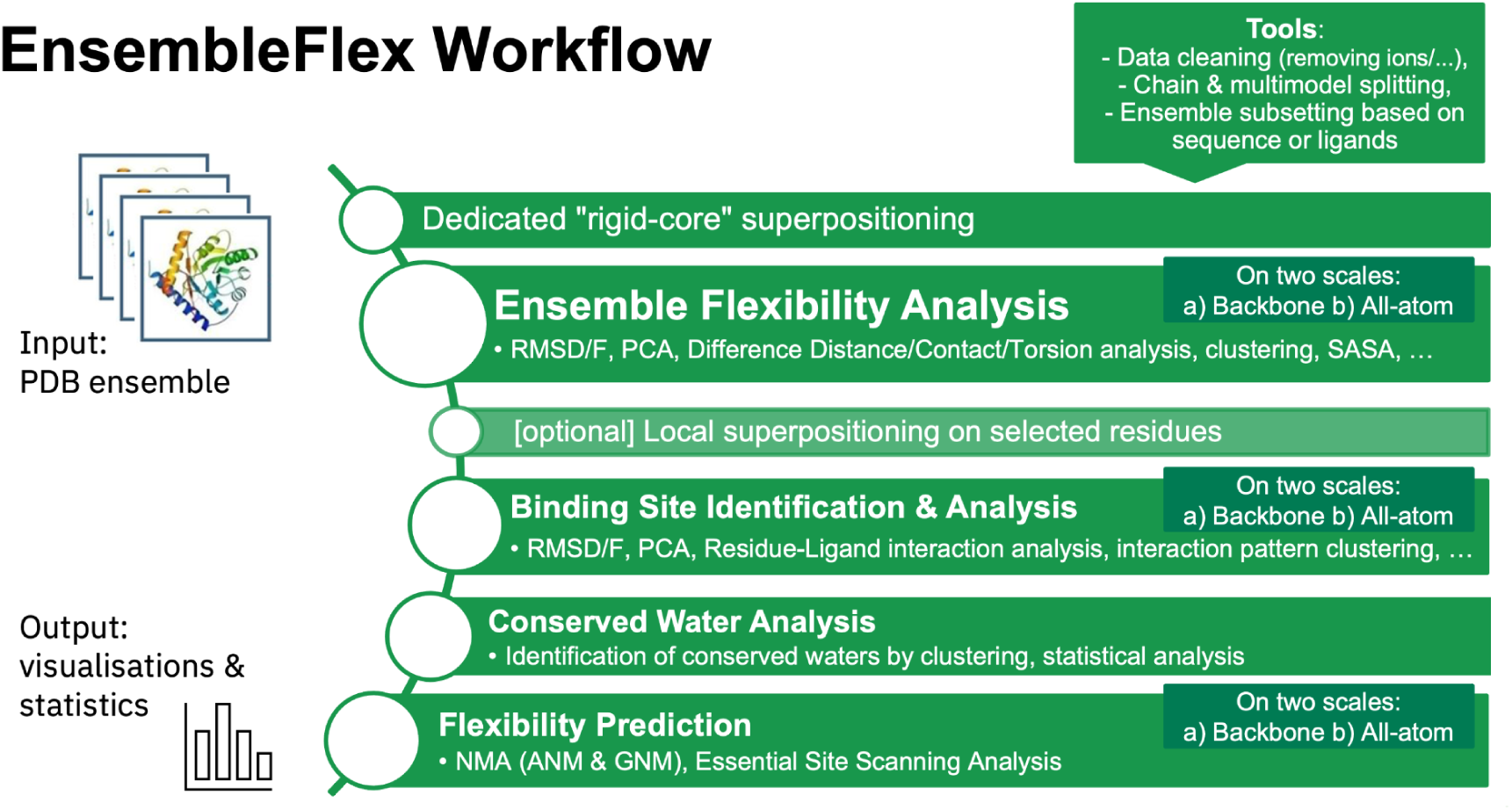
EnsembleFlex workflow overview

### Input and Output

The primary input for EnsembleFlex is a directory of PDB files representing a protein ensemble. The tool is designed to handle both uniform ensembles (e.g. NMR models, MD snapshots) and also heterogeneous ensembles (where atoms and/or residues differ across the ensemble, such as PDB ensembles), focusing on single-chain (monomeric) proteins. Outputs include various plots (histograms, heatmaps, etc.), modified PDB files augmented with analysed data (stored in the B-factor column), and PyMol scripts for visualisation. EnsembleFlex provides detailed information on flexibility measurements, ligand binding sites, and conserved waters, tailored for further analysis and integration into existing workflows. For example, RMSD and RMSF analyses, as well as Principal Component Analysis are provided to quantify structural deviations and fluctuations, offering insights into both global and local protein flexibility. The interface, as well as the provided user guide, include tooltips and an integrated explanation section (see Fig. S1), guiding users through each step of the analysis. In the interface the analytic (or predictive) calculations can be executed by simply clicking on “run” buttons. Subsequently, analysis results are presented in scrollable containers with tab navigation between thematic groups (see Fig. S2). The GUI provides a user-friendly presentation of key results, including interactive 3D structure visualizations enhanced with integrated data and animations. This facilitates and accelerates interpretation by directly highlighting key areas of the analysis on the protein structures.

### EnsembleFlex Analysis Workflow

#### Superposition Methods

Superposition is a critical first step in flexibility analysis, as the accuracy of subsequent analyses depends heavily on the alignment of structures.^26^ EnsembleFlex employs a superposition method based on rigid-core identification, specifically optimised for flexibility analysis, provided by the R package Bio3D.^4^ The implemented function uses the default settings and a maximal cumulative volume of 1 Å^3^ at which core positions are detailed. This approach contrasts with the traditional all-atom minimal RMSD method, providing improved accuracy in flexibility assessments. An alternative method, provided through the python package ProDy ^18^, is also available, and the user has the option to provide already structurally aligned conformations, skipping the superpositioning through EnsembleFlex.

#### Coordinate-Based and Difference-Based Flexibility Analysis Methods on two scales

EnsembleFlex integrates both backbone and all-atom flexibility analysis, using metrics like RMSD and RMSF to quantify structural variability. These analyses are complemented by difference distance, contact, torsion and Solvent Accessible Surface Area (SASA) analyses, which are essential for understanding protein dynamics at different scales.

RMSD measures the overall structural deviation between two conformations of a protein. It quantifies the average distance between the atoms of a target structure and a reference structure after optimal superimposition. A higher RMSD value indicates a greater structural difference between the two conformations. Nevertheless, RMSD is sensitive to larger changes in small numbers of residues, such as loops.

RMSF, on the other hand, measures the fluctuation of individual atoms or residues around their average positions within an ensemble or trajectory. It provides a residue-level measure of flexibility, highlighting the most mobile regions of the protein.

It’s important to note that RMSD and RMSF provide complementary information. While RMSD may capture global structural changes, RMSF highlights local flexibility. Additionally, the interpretation of these metrics can be influenced by factors like the alignment strategy, reference structure, and ensemble size.^26^

Further difference-based methods are implemented in EnsembleFlex that have the advantage of being independent from atomic coordinates and are therefore not affected by different superpositionings. Those are for example differences in torsion angles and differences in distance matrices.^27,28^ Differences between those metrics can be very indicative for flexibility within the structural ensembles. In various cases the use of torsion angles may be more sensitive and more robust compared to atomic coordinates for investigating different conformational states.^27^ Within EnsembleFlex torsion angle differences are only calculated on backbone atoms, excluding the side chains. On the other hand, all-atom calculations are performed using difference distance matrices. Additionally, a dedicated protocol called ensemble difference distance matrix (eDDM) analysis is implemented^20^, which enables users to assess significant differences between groups of structures.

The Solvent Accessible Surface Area (SASA) for each residue is calculated for the whole ensemble using the Biopython implementation ^21,29^ and the differences and standard deviation across the ensemble provide insights into local environment changes the residues are experiencing in different states.

#### Dimension Reduction and Clustering for visualisation and interpretation

Dimension reduction techniques enable the visualisation of high-dimensional data, such as atomic coordinates, in a more interpretable form, such as 2D plots.^26,11^ EnsembleFlex includes two dimension reduction methods, Principal Component Analysis (PCA) and Uniform Manifold Approximation and Projection (UMAP), which in combination allow to retain different aspects of the data’s underlying structure, due to their linear and non-linear nature, respectively.^30^

Clustering is a valuable tool for interpreting structural datasets, helping researchers identify patterns and areas of interest they may want to focus on in the next iteration of experiments. The desired number of clusters is set by the user with an initial default of 3. Based on visual inspection of the generated plots (2D scatter plots and dendrograms) the user may want to adjust this value and rerun the analysis. The summary cluster attribution plot gives a visual overview of how the structures have been partitioned based on all analysis results, including the usage of backbone and all-atom data and other processing results, such as RMSD, PCA, and UMAP. If a user struggles with deciding on the desired number of clusters, the user may want to investigate the automatically generated best cluster number estimation reports. EnsembleFlex generates automated best cluster number estimation reports using connectivity, Dunn index and Silhouette score as metrics on partitioning results generated through hierarchical, k-means, and k-medoids/PAM algorithms, implemented through the R package clValid.^22^ The cluster number estimation procedure is performed on data from backbone-based RMSD calculations, PCA on backbone coordinates, and UMAP on backbone coordinates, resulting in three reports.

#### Focus on Ligand Binding Sites

EnsembleFlex provides automated methods dedicated to the exploration of ligand binding sites. Binding site residues are identified based on a user-defined heavy atom distance cutoff (with a default of 3,5 Å) from ligand atoms. Note that all HETATM atoms (that have not been filtered beforehand) are considered as ligand atoms. Therefore, EnsembleFlex includes tools to remove ions and other non-relevant molecules, as well as an automated ligand detection procedure as help for filtering. The frequency of identification within the binding site is calculated for each residue and visualised in a histogram. For visual inspection the identified residues are labelled for each structure using the b-factor column of the given structure and saved as a pdb file. Further summary pdb files are provided containing the coordinates of the reference structure and in the b-factor column either whether a residue was identified as being part of the binding site in any structure of the ensemble (values of 0 or 1), or the calculated occurrence frequencies across the ensemble. The radius of gyration, as an approximation of size, is calculated for the identified binding site residues per structure and visualised in a histogram. Ligand-residue interactions are plotted as heatmap with pattern clustering to allow detection of ligand dependant characteristics, such as eventual sub-pockets.

#### Conserved Water Analysis

Conserved water molecules play a critical role in protein structure stabilisation, flexibility, and ligand binding. EnsembleFlex implements a streamlined conserved water analysis using a slightly modified version of the R package *vanddraabe*^24^, which is based on cluster analysis methods similar to those used in WatCH^31^ and PyWATER^32^. In this analysis, the tool identifies water molecules conserved across multiple crystallographic structures by clustering them based on their spatial proximity (oxygen atoms within 2.4 Å of each other are considered a cluster) after structural superposition. The degree of conservation is calculated as the number of water molecules present in a cluster divided by the total number of structures in the ensemble. Water molecules that are consistently located near protein atoms, in hydrophilic environments, and forming hydrogen bonds tend to be more conserved. This conserved water analysis is especially valuable in understanding the ligand-binding environment, as conserved water molecules often mediate key interactions between proteins and ligands. EnsembleFlex generates a detailed analysis of conserved waters, highlighting their role in the structural integrity and function of protein binding sites. The output includes 3D visualisations in PyMol and a statistical summary of conserved water clusters. By combining this analysis with ligand binding site analysis and flexibility metrics, EnsembleFlex offers a comprehensive picture that is highly relevant for drug design, where water-mediated interactions can be critical to ligand binding.

#### Data-Based and Predictive Methods

While EnsembleFlex primarily focuses on data-driven analysis, it also incorporates predictive methods based on elastic network models (ENM). These models, including Normal Mode Analysis (NMA), offer insights into protein dynamics that may not be fully captured by experimental data alone, but also have their own limitations, such as bias to the starting structure and conformational exploration limits. From the global backbone perspective EnsembleFlex provides two ENM based NMAs, Anisotropic Network Model (ANM) C-alpha Normal Mode Analysis (NMA) and Gaussian Network Model (GNM) C-alpha Normal Mode Analysis (NMA), both based on the reference structure (the first structure of the ensemble). In general, GNM is preferred when accurately predicting the distribution and magnitudes of residue fluctuations is the primary goal, while ANM is better suited for assessing the directionality and mechanisms of motions.^33^ The choice depends on whether the focus is on the amplitudes or the directions of the motions. From the focussed site-of-interest point of view EnsembleFlex provides an overall all-atom Normal Mode Analysis (aaNMA), implemented through Bio3D, and an Essential Site Scanning Analysis (ESSA), an ENM-based method that identifies residues that would significantly alter the protein’s global dynamics if bound to a ligand, implemented through ProDy.^34^ By combining computational predictions with experimental data, EnsembleFlex provides a more comprehensive understanding of protein flexibility.

## Results - Case Studies

The following four case studies contain two example protein ensembles, where all available structures of the given protein are downloaded from PDBe^35^ through their respective PDBe-KB “aggregated views” pages^36^, and two further example protein ensembles which represent group depositions downloaded from RCSB-PDB^37^ via their group ID.

### 1. Adenylate Kinase (E. coli, UniProt-ID: P69441)

#### Background and Interest

Adenylate kinase plays a pivotal role in cellular energy regulation by catalysing phosphate transfer between ATP and AMP. Its catalytic efficiency depends on its ability to transition between open and closed conformations, making its structural flexibility critical to understanding its mechanism.^38,39^

#### Flexibility Analysis Results

The ensemble from the PDBe-KB aggregated views page (www.ebi.ac.uk/pdbe/pdbe-kb/proteins/P69441) comprises 21 PDB structures with 39 monomers, which reveal two distinct conformational states across all assessments except UMAP clustering (see cluster attribution plot in Fig. 2A). This discrepancy in UMAP highlights how certain dimensional reduction techniques may emphasise different aspects of the data. The PCA results are particularly revealing: PC1 contributes the most to distinguishing the two states, with a remarkable proportion of variance of 96.92% (see Fig. 2C). The RMSD analysis (heatmap and cluster dendrogram, Fig. 2D), as well as the other analysis confirm the exact same bipartition. The PC1 residue contribution plot pinpoints specific regions critical to this transition. RMSF plots further highlight these key residues showing differences in flexibility between the two conformational states (see Fig. 2E).

**Fig. 2:**
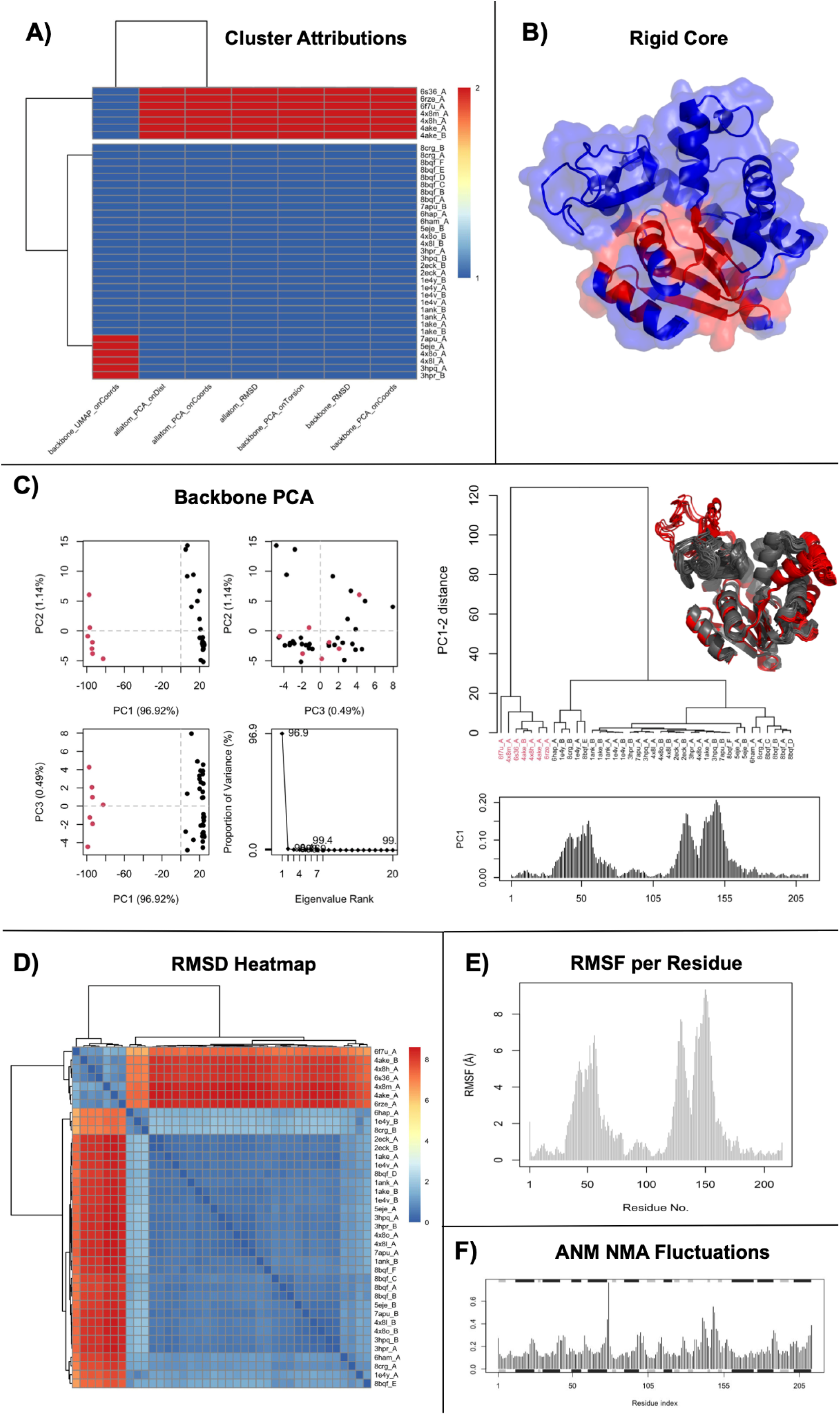
Adenylate kinase analysis: A) Cluster attributions for the whole dataset by all seven methods; B) Identified rigid-core residues, which were used for superpositioning colored in red on reference structure; C) PCA with first three PCs plotted against each other and proportion of variance of PC1-20, and PC1-2 cluster dendrogram with all superimposed structures colored by group, and PC1 residue contribution; D) Pairwise RMSD heatmap; E) Backbone RMSFs per residue; F) ANM NMA fluctuations

The analysis also leverages a predictive normal mode analysis (NMA) approach. Here, elastic network models (ANM fluctuations shown in Fig. 2F) suggest a less pronounced flexibility variation across different protein regions compared to the experimental data-driven methods. This highlights the complementarity between predictive and data-based methods: while predictive models provide generalised insights, the data-driven analysis captures subtle, functionally relevant dynamics specific to the experimental dataset.

#### Insights and Implications

These results emphasise the importance of combining data-driven and predictive approaches to gain a comprehensive view of flexibility. The residue-specific flexibility differences identified through PCA and RMSF can guide further functional studies and drug design efforts targeting specific conformational states.

### 2. Hexokinase-1 (UniProt-ID: P19367)

#### Background and Interest

Hexokinase-1 initiates glucose metabolism by converting glucose into glucose-6-phosphate. This enzyme undergoes large-scale domain movements, including hinge bending and rotational shifts, to accommodate substrate binding and catalysis. Understanding these dynamics is vital for comprehending its functional mechanism.^40,41^

#### Flexibility Analysis Results

The ensemble from the PDBe-KB aggregated views page (www.ebi.ac.uk/pdbe/pdbe-kb/proteins/P19367) includes 10 PDB structures and 17 monomers. On the PDBe aggregated views page, the superposition is performed globally, resulting in a “blurred” ensemble with conformational variation seemingly all over the protein structure, lacking a clear distinction of where the actual conformational changes occur and which parts can be considered as rigid. In contrast, EnsembleFlex employs a dedicated rigid-core superposition method, which provides a much clearer vision of the structural dynamics by aligning the ensemble based on the most stable regions of the protein. The subsequent RMSF analysis on two scales - backbone only and all-atom - provides a detailed view on dynamics (see Fig. 3A/B). The backbone RMSF analysis clearly identifies the two domain movement, whereas the all-atom RMSF highlights additional flexibility of side chains within the two domains. PCA results reveal large-scale domain movements: PC1 captures the hinge motion, while PC2 reflects domain rotation as shown by the extrapolation animations along the PCs (see Fig.3C). These findings emphasise the importance of the flexibility analysis workflow, especially the use of rigid-core superposition methods, which avoid artefacts and a blurred view introduced by traditional all-atom RMSD superposition.

**Fig. 3:**
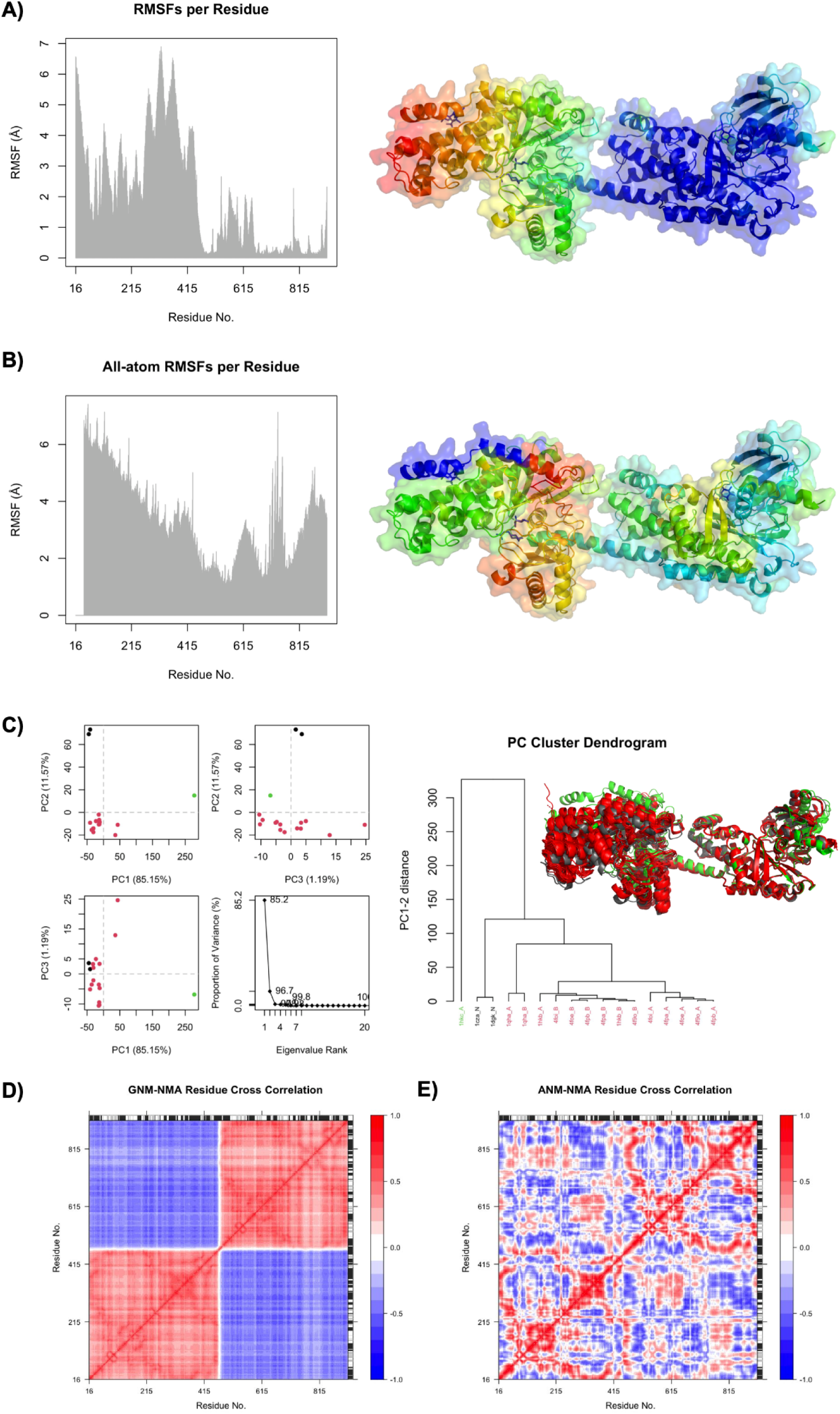
Hexokinase-1 analysis: A) Backbone RMSF along the sequence and colour-coded on the reference structure (scheme rainbow: blue-green-yellow-red); B) All-atom RMSF along the sequence and colour-coded on the reference structure (scheme rainbow: blue-green-yellow-red); C) PCA with first three PCs plotted against each other and proportion of variance of PC1-20, and PC1-2 cluster dendrogram with superimposed conformations colored by PC1-2 cluster; D) GNM and E) ANM residue cross correlations.

Predictive NMA analysis complements these findings. The GNM-based model captures a two-domain motion that aligns with the data-driven PCA results (see Fig. 3D). However, the ANM-based model does not highlight this movement as explicitly, but instead puts more emphasis on within domain movements (see Fig. 3E), underscoring the limitations of relying solely on predictive methods. This reinforces the importance of integrating multiple approaches for a robust analysis of structural flexibility.

#### Insights and Implications

By identifying the distinct contributions of hinge and rotational motions, the analysis provides detailed insights into Hexokinase-1’s functional mechanics. These results can inform future studies, particularly those exploring allosteric regulation or targeted inhibition strategies.

### 3. Interleukin-1 Beta Fragment Screen (Group-ID: G_1002139)

#### Background and Interest

Interleukin-1 beta is a cytokine central to inflammation and immune response.^42^ The ensemble (registered under the group ID G_1002139, accessible through the RCSB-PDB site at www.rcsb.org/groups/summary/entry/G_1002139), derived from a fragment screening experiment^43^, is a rich dataset for studying flexibility and ligand interactions, crucial for drug discovery efforts targeting this protein and its interactions.

#### Flexibility Analysis Results

The ensemble consists of 22 monomers and identifies two distinct conformational states, primarily driven by loop movements. Flexibility assessments, including backbone-only and all-atom methods, converge on these findings. Loop movement regions, detected through RMSF and PCA residue contribution plots, highlight flexible residues critical for function and ligand interaction (see Fig. 4).

**Fig. 4:**
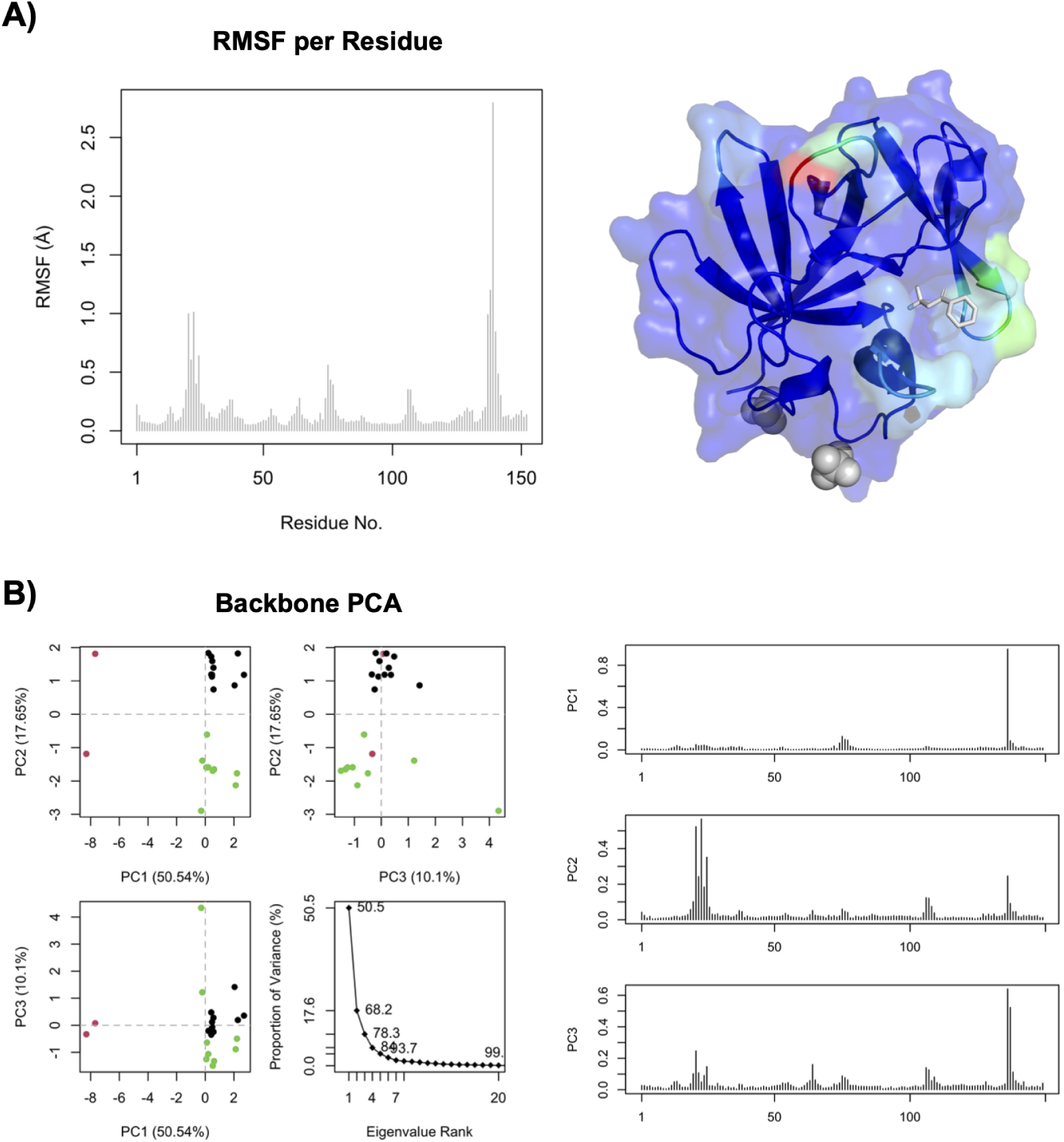
Interleukin-1 beta fragment screen analysis: A) Backbone RMSF per residue as barplot and colour-coded on the reference structure (scheme rainbow: blue-green-yellow-red); B) Backbone PCA with first three PCs plotted against each other and proportion of variance of PC1-20, and residue contributions for PC1-3;

Ligand-induced dynamics are explored through a heatmap of ligand-protein interactions (see Fig. 5A). This analysis groups ligands based on shared binding patterns, revealing clusters of structurally and functionally similar ligands. A quantification of frequency of involvement in ligand binding is provided as a ranked barplot and values are color-coded onto the reference structure (see Fig. 5B), which enables a focused binding site assessment. An all-atom mean RMSF histogram of binding site residues (see Fig. 5C) provides further insight, identifying particularly flexible residues. These residues could influence binding affinity and stability, guiding prioritisation of ligand candidates for further development.

**Fig 5:**
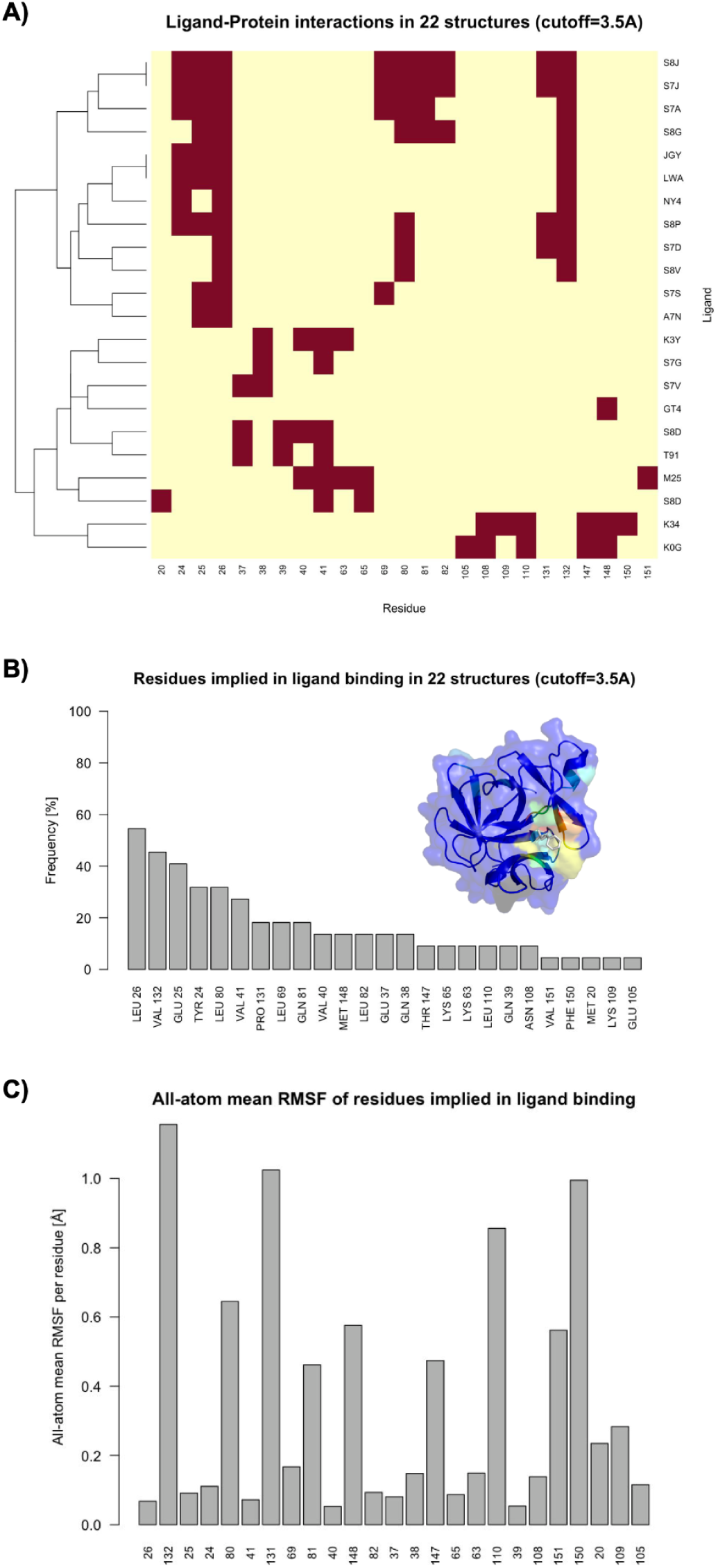
Interleukin-1 beta fragment screen analysis: Residues involved in ligand binding A) as interaction matrix, B) as quantifying barplot ordered by frequency of involvement, with reference structure coloured by frequency of involvement (scheme rainbow: blue-green-yellow-red), C) and their respective all-atom mean RMSF, ordered as in B.

Complementary predictive analysis using Essential Site Scanning Analysis (ESSA) identifies residues in the binding site that may significantly influence global protein dynamics. In the case of Interleukin-1 beta ensemble several binding site residues are above the line, and particularly residue Q39 stands out (see Fig. 6A), which at the same time shows very little flexibility in the RMSF plot (Fig. 5C), suggesting that perturbing the conformation of this residue may have a larger impact. Color-coding the ESSA z-scores on the reference structure (shown in Fig. 6B) helps identify areas that may be more or less prone to dynamic perturbations. Conserved water analysis identifies water molecules critical to ligand binding, offering opportunities for fragment expansion and optimization. The global water analysis plot (Fig. 6C) gives an overview on ensemble statistics and the respective 3D visualisation in PyMol (Fig. 6D) directly provides the three-dimensional action points.

**Fig. 6:**
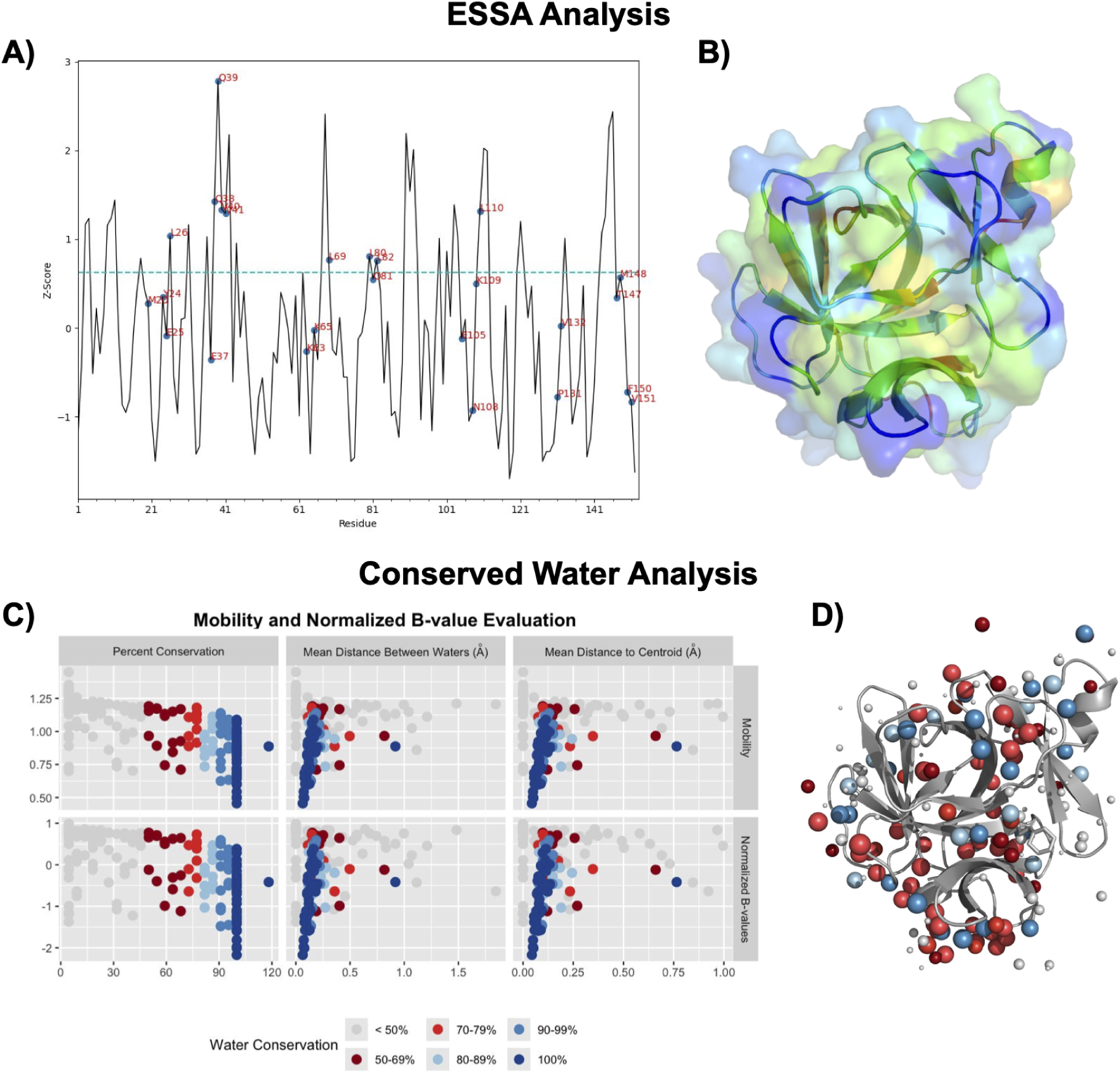
Interleukin-1 beta fragment screen - ESSA analysis: A) GNM Z-Scores of ESSA plotted against the protein sequence with highlighted binding site residues; B) Reference structure coloured by GNM Z-Score values of ESSA (scheme rainbow: blue-green-yellow-red); Conserved Water Analysis: C) Global mobility and normalized B-value evaluation D) Conserved waters as spheres with size based on conservation q (vdw=((q/10) / (5 * 3.14**2))**0.5 * 25), colored as in A together with the reference structure in grey.

#### Insights and Implications

The combined insights from flexibility, binding site dynamics, and conserved water analysis provide a comprehensive framework for ligand prioritisation and optimization. The integration of data-driven and predictive methods enhances the understanding of flexibility and ligand binding, making the generated data invaluable for drug design.

### 4. Covid Moonshot Project on SARS-CoV-2 Main Protease (Group-ID: G_1002272)

#### Background and Interest

The SARS-CoV-2 main protease is essential for viral replication and a prime antiviral target. This ensemble, a result of the collaborative Covid Moonshot Project^44^, currently represents the largest group deposition registered in the RCSB-PDB (Group-ID: G_1002272, accessible at www.rcsb.org/groups/summary/entry/G_1002272), providing a unique opportunity to study large-scale flexibility in a critical protein.

#### Flexibility Analysis Results

The ensemble comprises 466 structures and 693 monomers, revealing three distinct conformational groups consistently across both scale analyses (backbone-only and all-atom, see Fig. 7). RMSD, PCA, and distance matrix analyses all align, demonstrating the robustness of the flexibility analysis workflow, even for such a large dataset. Dimension reduction methods, dendrograms and heatmaps, effectively visualise the clustering of these groups, making the data interpretable despite its scale. When simply looking at the superimposed structures (Fig. 7B), the clustering into three distinct groups would not be obvious, highlighting the need for quantification with computational tools.

**Fig. 7:**
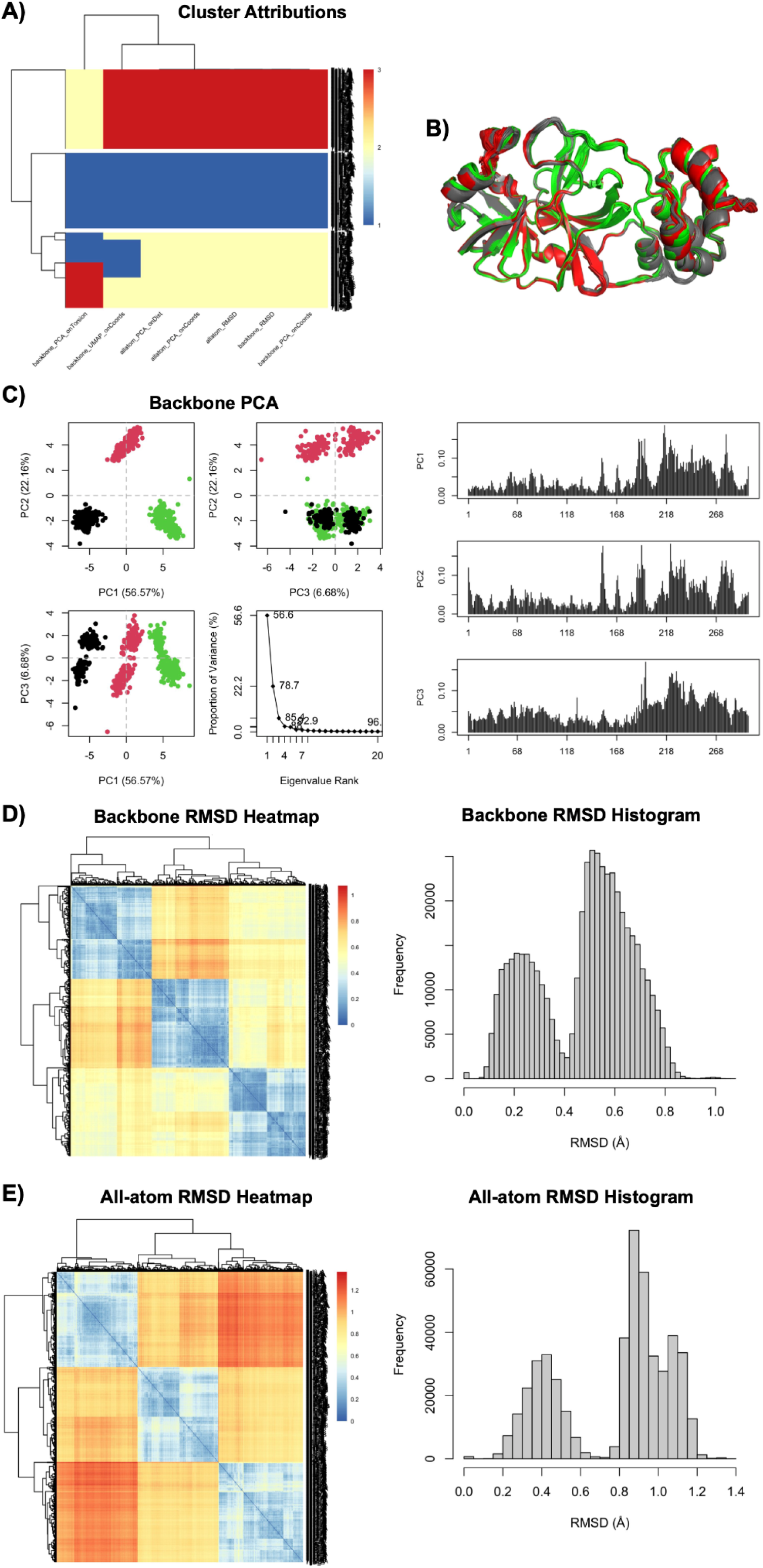
Covid Moonshot Project analysis: A) Cluster attributions for the whole dataset by all seven methods; B) Superimposed structures colored by backbone PCA cluster attribution; C) Backbone PCA with first three PCs plotted against each other and proportion of variance of PC1-20, and residue contributions for PC1-3; D) Backbone RMSD heatmap and histogram; E) All-atom RMSD heatmap and histogram.

The results emphasise the scalability and reliability of EnsembleFlex in handling extensive datasets. This capability is crucial for addressing the challenges posed by large-scale drug discovery projects, where datasets often exceed hundreds of structures.

#### Insights and Implications

By identifying distinct conformational groups, the analysis provides a foundation for studying functional dynamics and designing inhibitors that target specific conformational states. The robustness of the methods across diverse scales demonstrates EnsembleFlex’s versatility.

## Discussion

### Advantages of EnsembleFlex

EnsembleFlex combines methodological precision with user-friendly design to significantly enhance the analysis of protein structure ensembles. Its specialised superposition method, tailored for flexibility analysis, ensures accurate alignment of structures and avoids artefacts introduced by standard all-atom RMSD methods. The importance of this approach is evident in the Hexokinase-1 ensemble, where rigid-core superposition accurately captures domain movements such as hinge bending and rotational shifts - critical for understanding functional mechanics. Similarly, in Adenylate Kinase, the accurate superposition enabled the detection of distinct conformational states, providing insights into the residues driving these transitions.

The integration of clustering and dimension reduction techniques further aids in making sense of complex datasets, especially large-scale ensembles like the screening on SARS-CoV-2 main protease. Dimension reduction methods effectively visualised the three conformational groups identified, ensuring clarity even in datasets comprising hundreds of monomers. The use of multiple descriptors, such as coordinates, distances, and torsion angles in combination with dimension reduction techniques, such as PCA and UMAP, allows researchers to retain and interpret different facets of the underlying data, making EnsembleFlex a valuable tool for experimental scientists.

The tool’s dual-scale flexibility analysis, encompassing both backbone and side-chain dynamics, provides a holistic view of protein behaviour. For instance, in the Interleukin-1 beta screening ensemble, the combination of backbone and all-atom analyses highlighted flexible residues in binding sites that influence ligand affinity. Integrating predictive tools like GNM-based normal mode analysis complements these data-driven insights, filling gaps in experimental datasets. This is particularly evident in the Adenylate Kinase ensemble, where predictive NMA revealed generalised flexibility patterns, enhancing the understanding provided by data-based analyses.

Moreover, EnsembleFlex offers advanced capabilities, such as conserved water analysis, which are crucial in drug discovery. In the Interleukin-1 beta screening ensemble, conserved water analysis identifies opportunities for fragment expansion, providing actionable insights for fragment-based ligand design. The ligand-protein interaction heatmap and RMSF-based residue flexibility analysis further facilitate the prioritisation of ligand candidates.

In high-throughput experimental contexts, such as fragment screening and automated crystallography workflows at facilities like the HTX Lab at EMBL Grenoble ^13,45^, EnsembleFlex serves as a vital tool for analyzing large datasets. By providing accessible and streamlined analysis capabilities, it enhances the utility of structural data generated from such platforms.

### Applications Across Domains

The wide applicability of EnsembleFlex spans drug design, protein engineering, and fundamental structural biology. For example:

- In **drug design**, flexibility analysis reveals how conformational changes influence ligand binding, as seen in the Interleukin-1 beta screening example. By identifying flexible and conserved regions, researchers can prioritise ligand candidates and design inhibitors that target specific conformational states.
- In **protein engineering**, the understanding of hinge motions and domain rotations, such as in Hexokinase-1, aids in designing proteins with enhanced stability or specific functional properties.^46^
- In **structural biology**, tools like EnsembleFlex streamline the analysis of dynamic ensembles, making large-scale studies like the Covid Moonshot project on the SARS-CoV-2 main protease manageable and insightful.

In research projects where flexibility plays a critical role combining different analysis tools is essential to fully capture the protein’s dynamic behaviour. For example, backbone flexibility could reveal how a protein changes its conformation upon ligand binding or as a result of protein-protein interactions, while side-chain flexibility highlights adjustments in the active site that directly affect binding affinity. Combining these results with conserved water analysis can provide additional insights into how water molecules stabilise the ligand-bound state.

The user-friendly interface and coding-optional design lower the barrier to entry, enabling a broader audience to perform advanced flexibility analyses without extensive computational expertise.

### Limitations and Future Work

Despite its strengths, EnsembleFlex has limitations. The tool is currently tailored for monomeric proteins, making the analysis of multi-chain systems challenging. Expanding its capabilities to handle multi-chain complexes would enhance its utility in analysing large assemblies and multimeric enzymes. Additionally, while the predictive methods integrated into the tool are robust, incorporating advanced approaches like deep learning-based flexibility prediction could further improve accuracy and complementarity with experimental data. Enhancing interactive visualisation features could make the tool even more intuitive for users.

## Conclusion

### Summary

EnsembleFlex represents a significant advancement in protein flexibility analysis by integrating specialised superposition, clustering, and dimension reduction methods into a single platform. Its dual-scale approach bridges the gap between backbone-only analyses and the more nuanced all-atom dynamics critical for understanding ligand interactions and functional flexibility. By combining predictive models with data-driven methods, EnsembleFlex provides a comprehensive perspective on protein dynamics.

The case studies underscore the tool’s versatility and impact:

● The Adenylate Kinase analysis demonstrates the tool’s ability to detect distinct conformational states and critical residues.
● The Hexokinase-1 analysis showcases its capacity to resolve large-scale domain movements using rigid-core superposition.
● The Interleukin-1 beta screening analysis highlights its application in ligand screening and fragment prioritisation, supported by insights into conserved water interactions.
● The Covid Moonshot dataset analysis exemplifies its scalability, handling extensive datasets while maintaining analytical precision.

### Impact

EnsembleFlex’s user-friendly design and comprehensive analysis suite make it an indispensable tool for researchers across structural biology, drug design, and protein engineering. By offering a comprehensive suite of tools for protein ensemble analysis, EnsembleFlex empowers researchers to uncover deeper insights into protein structure-function relationships. Its user-friendly design ensures accessibility for experimental and computational scientists alike, fostering its adoption across various fields. Whether addressing large-scale projects, such as the Covid Moonshot Project on SARS-CoV-2 main protease, or providing critical ligand-binding insights in cases like the fragment screening on Interleukin-1 beta, EnsembleFlex stands out as an indispensable tool for advancing molecular design and therapeutic innovation. By extending the analytical capabilities of high-throughput facilities such as the HTX Lab at EMBL Grenoble, EnsembleFlex bridges the gap between experimental data generation and its interpretation, promoting innovation in molecular design and structural biology. EnsembleFlex’s ability to integrate experimental data with predictive modelling in a scalable and accessible platform sets a new benchmark for protein flexibility analysis. By bridging gaps in our understanding of protein dynamics, it holds the potential to accelerate research in drug design, protein engineering, and beyond, enhancing the quality and impact of scientific discoveries.

## Acknowledgements

This work was supported by the ARISE Fellowship from the European Union’s Horizon 2020 research and innovation program under the Marie Skłodowska-Curie grant agreement No 945405. It also benefited from the Fragment Screen project (No 101094131), funded by the Horizon 2020 program. The authors thank contributors who assisted in the testing of EnsembleFlex.

## Appendices

### Supplementary Data

Code/tool availability: https://gitlab.ebi.ac.uk/melanie/ensembleflex

Reference to the Bio.tools registry: https://bio.tools/ensembleflex

Installation instructions are provided in the README.md file, as well as the user_guide.md file.

**Fig S1.:**
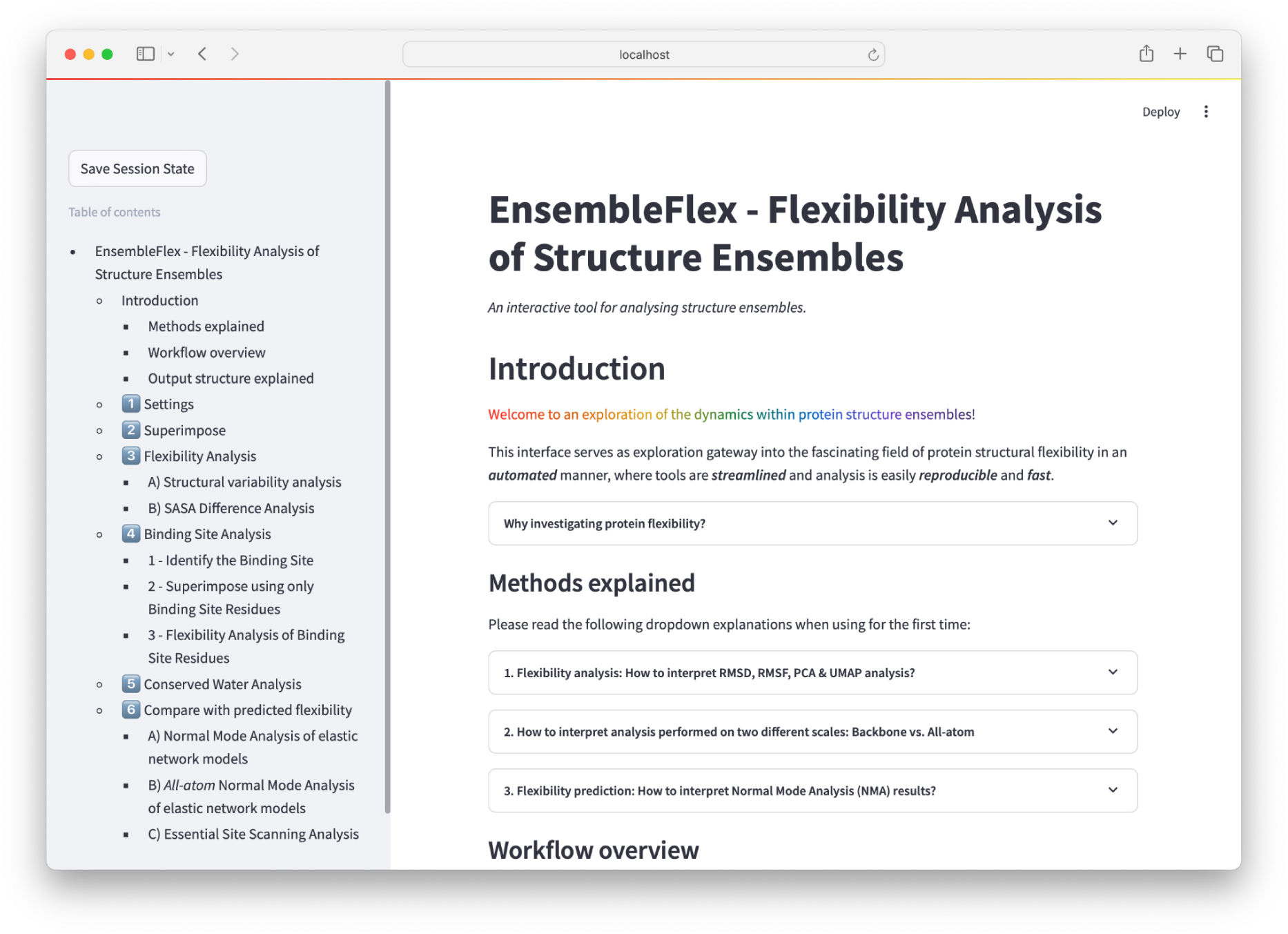
GUI screenshot in browser. The GUI is a scrollable page with a side panel displaying the contents for ease of navigation.

**Fig S2.:**
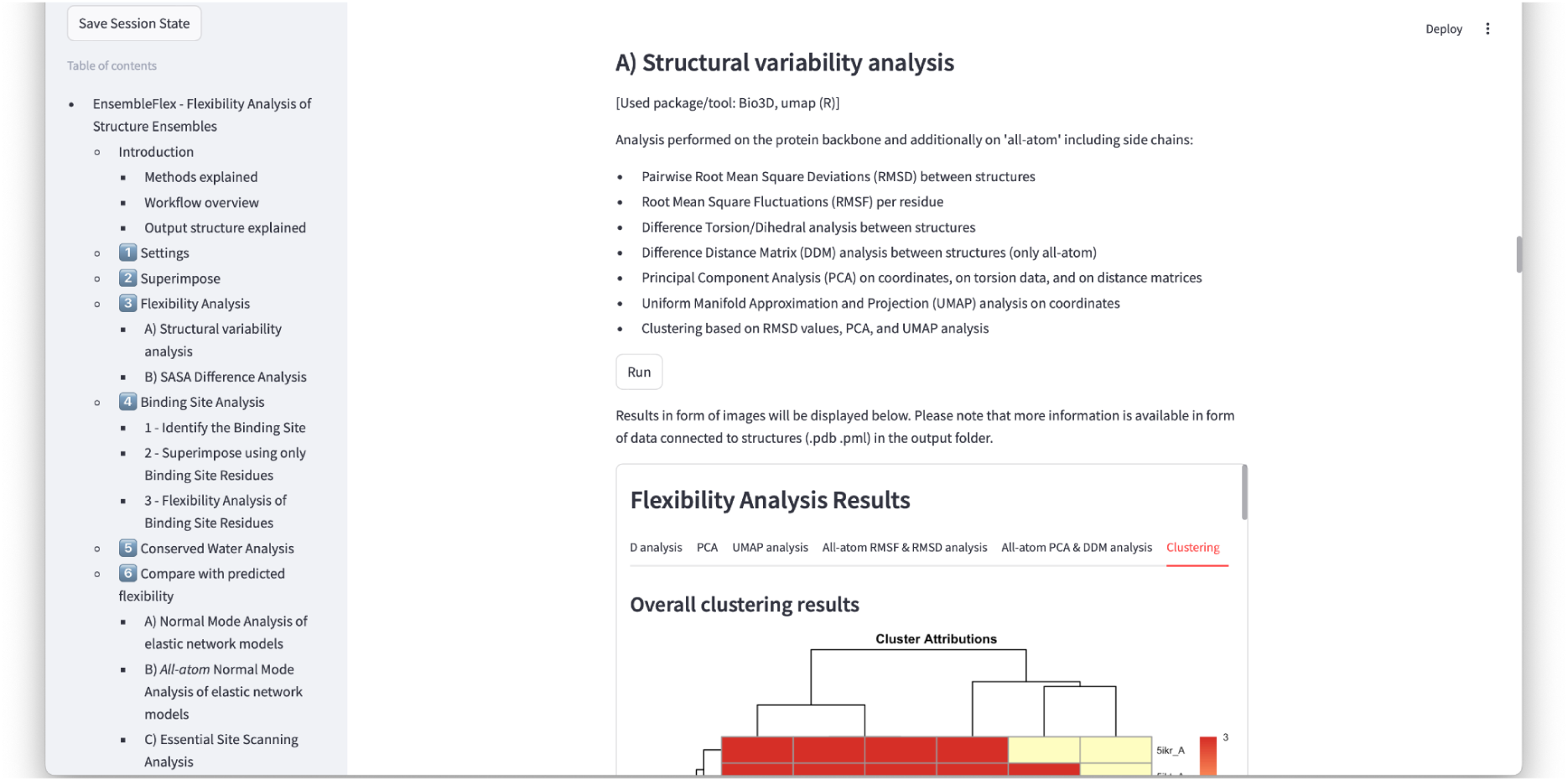
GUI screenshot showing the main flexibility analysis section with its execution button “Run” and the scrollable result container with its thematic tabs.

## Notes

https://gitlab.ebi.ac.uk/melanie/ensembleflex

https://bio.tools/ensembleflex

### Competing Interest Statement

The authors have declared no competing interest.

